# Temporal dynamics of color processing measured using a continuous tracking task

**DOI:** 10.1101/2024.03.01.582975

**Authors:** Michael A. Barnett, Benjamin M. Chin, Geoffrey K. Aguirre, Johannes Burge, David H. Brainard

## Abstract

We characterized the temporal dynamics of color processing using a continuous tracking paradigm by estimating temporal impulse response functions associated with tracking chromatic Gabor patches. We measured how the lag of these functions changes as a function of chromatic direction and contrast for stimuli in the LS cone contrast plane. In the same set of subjects, we also measured detection thresholds for stimuli with matched spatial, temporal, and chromatic properties. We created a model of tracking and detection performance to test if a common representation of chromatic contrast accounts for both measures. The model summarizes the effect of chromatic contrast over different chromatic directions through elliptical isoresponse contours, the shapes of which are contrast independent. The fitted elliptical isoresponse contours have essentially the same orientation in the detection and tracking tasks. For the tracking task, however, there is a striking reduction in sensitivity to signals originating in the S cones. The results are consistent with common chromatic mechanisms mediating performance on the two tasks, but with task-dependent relative weighting of signals from L and S cones.

## Introduction

In human vision the retinal image is encoded by the L, M, and S cones (Brainard & Stockman, 2010). Subsequent stages of processing combine the signals from the three classes of cones to create, broadly speaking, three post-receptoral mechanisms: two cone-opponent mechanisms and a luminance mechanism. The cone-opponent mechanisms represent the differences between cone signals [S-(L+M) and L-M] while the luminance mechanism represents an additive combination (L+M) (Stockman & Brainard, 2010). The physiological basis of these mechanisms begins in the retina, with cone-opponent responses observed in retinal ganglion cells as well as in their targets in the lateral geniculate nucleus (e.g., DeValois, Abramov, & Jacobs, 1966; Derrington, Krauskopf, & Lennie, 1984; see Lennie & Movshon, 2005; Shevell & Martin, 2017). The sensitivity of each of the three cone-opponent post-receptoral mechanisms varies in a distinct manner with spatial and temporal frequency (e.g., de Lange, 1958; Kelly, 1975; Mullen, 1985; Sekiguchi, Williams, & Brainard, 1993; Poirson & Wandell, 1996; Metha & Mullen, 1996).

Models based on three post-receptoral mechanisms are able to account for aspects of the detection and discrimination of colored patterns (e.g., Guth, Massof, & Benzschawel, 1980; Krauskopf, Williams, & Heeley, 1982; Poirson, Wandell, Varner, & Brainard, 1990; Poirson & Wandell, 1990b; Poirson & Wandell, 1990a; Guth, 1991; Poirson & Wandell, 1993; Poirson & Wandell, 1996; Knoblauch & Maloney, 1996), although deviations from model predictions have suggest the presence of additional mechanisms, perhaps at cortical sites (e.g., Krauskopf, Williams, Mandler, & Brown, 1986; Gegenfurtner & Kiper, 1992; Eskew, Wang, & Richters, 2004; Hansen & Gegenfurtner, 2013; see Gegenfurtner, 2003; Eskew, 2009; Stockman & Brainard, 2010).

Although S cones have temporal dynamics similar to those of L and M cones (Schnapf, Nunn, Meister, & Baylor, 1990; Stockman, MacLeod, & Lebrun, 1993), human detection sensitivity declines more rapidly with temporal frequency for stimuli that isolate S cones than for those that excite the L and M cones (Stockman, MacLeod, & DePriest, 1991). Similarly, reaction times to detect S-cone mediated stimuli are longer than for stimuli detected by L and M cones (Smithson & Mollon, 2004; McKeefry, Parry, & Murray, 2003). Mollon and Krauskopf (1973) measured reaction time as a function of background illuminance for 430, 500, and 650 nm stimuli and found reaction times of ∼300-400 ms with the highest latencies associated with the 430 nm stimuli. Using a two-pulse detection method, Shinomori and Werner (2008) measured temporal sensitivity for increments and decrements of S-cone isolating modulations and derived lags of 50– 70 and 100–120 ms, respectively. Consistent with these behavioral observations, studies using fMRI reveal that the response in early human visual cortex to S-cone stimuli is attenuated more rapidly with temporal frequency than for L- and M-cone stimuli processing for such stimuli (Engel, Zhang, & Wandell, 1997; Liu & Wandell, 2005; Spitschan, Datta, Stern, Brainard, & Aguirre, 2016; Gentile, Spitschan, Taskin, Bock, & Aguirre, 2024). And, the response of at least some single units to signals from the S cones is delayed relative to their responses to signals from the L and M cones (Cottaris & De Valois, 1998). Motion perception is also impoverished for stimuli detected only by the S cones (Cavanagh, MacLeod, & Anstis, 1987; Dougherty, Press, & Wandell, 1999). Per unit contrast, the strength of S-cone input to motion-selective cortex is small relative to L and M cone contributions (Seidemann, Poirson, Wandell, & Newsome, 1999; Wandell et al., 1999).

Visual perception supports tasks beyond stimulus detection and discrimination, including visually-guided behavior. For this reason, there has been recent interest in investigating the relation between perceptual-motor performance and the mechanisms of early visual processing (Bonnen, Burge, Yates, Pillow, & Cormack, 2015; Bonnen, Huk, & Cormack, 2017; Chin & Burge, 2022; Straub & Rothkopf, 2022; Burge & Cormack, 2023). A compelling approach—continuous target-tracking psychophysics (Bonnen, Burge, Yates, Pillow, & Cormack, 2015)—measures the ability of a subject to use an on-screen cursor to track a target undergoing a random walk (Brownian motion). The resulting data provides an efficient way to estimate the temporal impulse response function (tIRF) of the visual-motor system that is associated with the tracking behavior (Figure 1).

**Figure 1:**
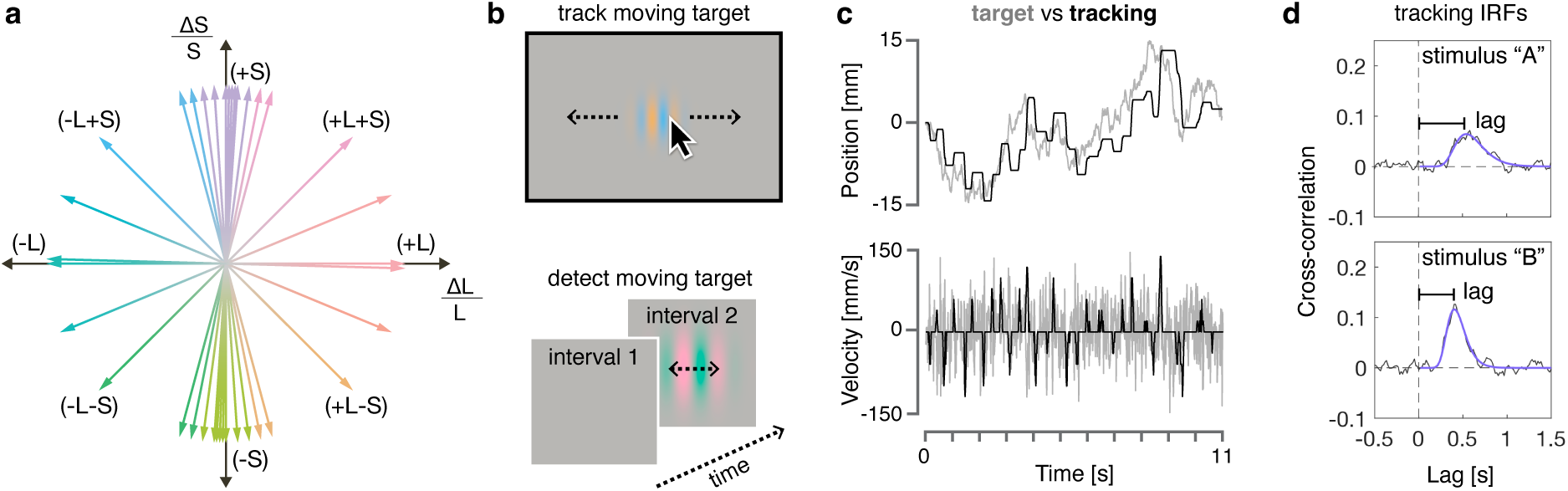
Experimental overview. (a) The LS cone contrast plane. The x-axis shows L-cone contrast and the y-axis S-cone contrast. Stimulus modulations around a background are represented as vectors in this plane, with the direction providing the relative strength of the L- and S-cone contrasts and the magnitude providing the overall contrast of the modulation. Indeed, we specify stimuli by their angular direction in the LS contrast plane, and their overall contrast by their vector length in this plane. A set of example directions are shown, with the color of each limb providing the approximate appearance of that portion of the modulation. (b) Example stimuli. In the tracking task, subjects tracked the position of a horizontally moving color Gabor modulation. In the detection task, subjects had to indicate which of two temporal intervals contained a horizontally moving color Gabor modulation. (c) The top panel shows position traces for an example tracking run in the experiment. The grey line shows the position of the target center as a function of time. The black line shows the subject’s cursor position as a function of time. The bottom panel show the velocities for the example data in the panel above. The grey line shows the velocity of the target as a function of time. The black like shows the cursor velocity as a function of time. (d) Example tracking impulse response functions (tIRFs). The grey line shows the cross-correlation between the stimulus and cursor velocities. This cross-correlation function is fit with a log-Gaussian function, and the peak of the function provides the tracking lag. Two example tIRFs are given for stimuli that evoke longer (stimulus “A”) and shorter (stimulus “B”) lags.

Here, we examined how cone signals are combined to support stimulus tracking. In two separate experiments we measured the ability of participants i) to track and ii) to detect targets that were matched in spatial, temporal, and chromatic properties. Figure 1 provides an overview of the stimuli and experimental paradigm. In both experiments, we characterized how performance varied for stimuli modulated in different directions in the L- and S-cone contrast plane. More specifically, we measured tracking lag, taken as the time to peak of the estimated tIRF associated each stimulus modulation (Figure 1d). We then employed a model-based approach to understand how the visual system integrates chromatic information to perform tracking and detection. The results characterize how any combination of L and S cone contrast contributes to performance on these tasks. We assess whether the data are consistent with the hypothesis that a common set of cone-opponent mechanisms mediate performance on both tasks. The data indicate a deficit in the ability of S-cones to support tracking, relative to detection. This deficit, however, is consistent with a common set of underlying cone-opponent mechanisms mediating the initial stimulus encoding for both tasks but with task-dependent readouts of the signals from these mechanisms.

## Results

### Experiment 1: Chromatic contrast and tracking lag

We first examined the basic relationship between the contrast of a stimulus and tracking lag. Figure 2 presents tracking lag as a function of contrast for Subject 2, grouped by the chromatic direction of the stimuli. Overall, we observed that tracking lag decreased as stimulus contrast increased. This was observed in all subjects and for all chromatic directions (see Figure S1 for data from Subjects 1 and 3).

**Figure 2:**
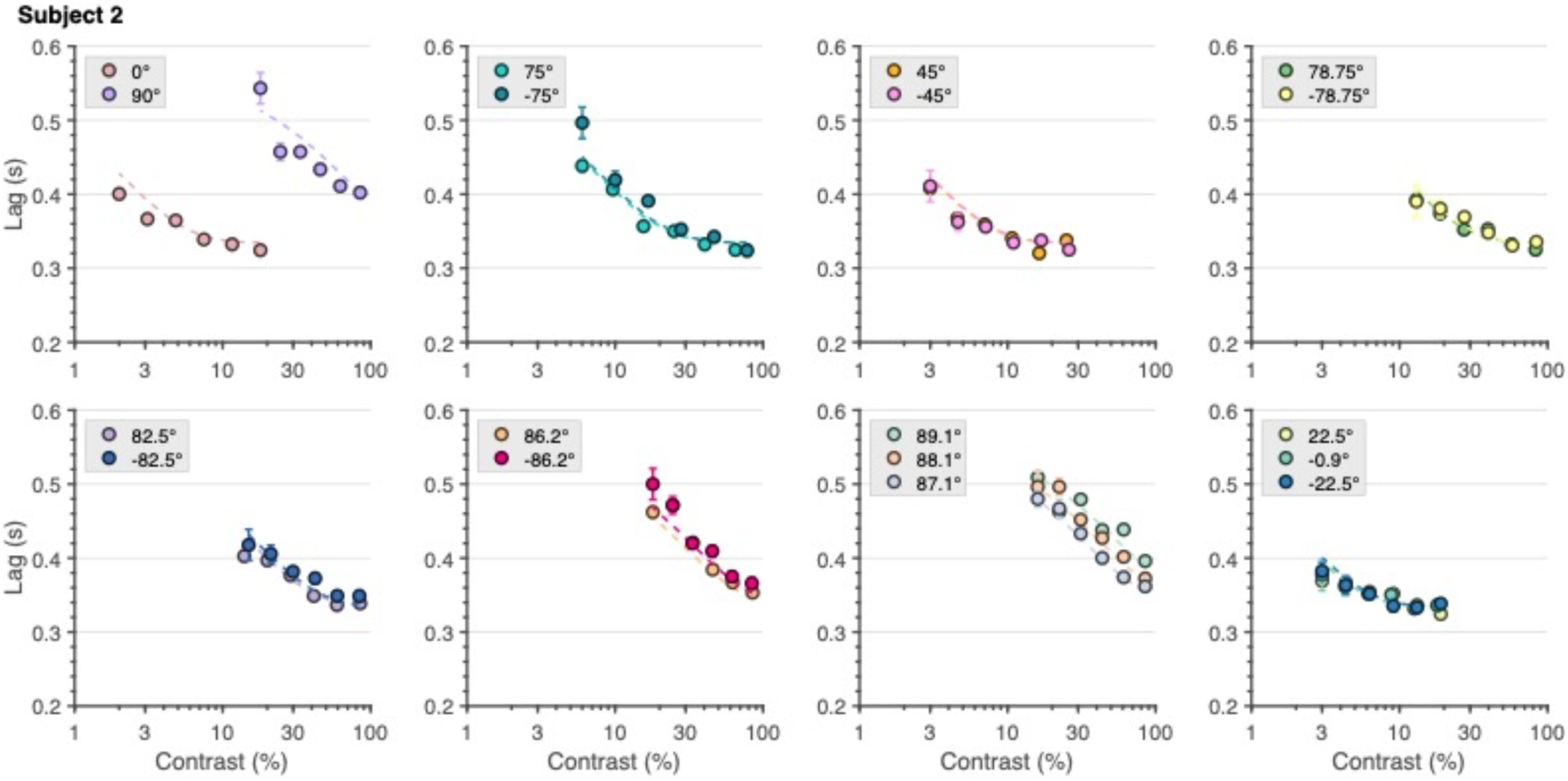
Lag Versus Contrast for Subject 2. Tracking lag as a function of modulation contrast separately for each chromatic direction. The closed circles indicate the lags of the tIRFs and are grouped into their corresponding chromatic direction by plot colors. The chromatic direction angles are displayed in the legend of each panel. The error bars on each lag estimate are the SEMs found via a bootstrap procedure (that is, the standard deviations of the bootstrapped parameter estimates). The dashed curves in each panel are the lag predictions of the color tracking model (CTM; see the following sections). The line colors lines are matched to the color of the corresponding symbols.

While tracking lag decreases with increases in contrast, the rate of decrease and minimum lag differed across chromatic directions. For example, the lag for tracking a 20% contrast L-cone isolating (0°) stimulus was roughly 325 ms, while 70% contrast was needed to achieve this low tracking latency for ±75° directions. Notably, tracking an S-cone isolating (90°) stimulus was associated with the longest lag values in all three subjects. This observation is broadly consistent with the prior literature on S-cone mediated temporal and motion processing (see Introduction).

With sufficiently high contrast, lag tended to asymptote at the same value for chromatic directions that incorporated some amount of L-cone contrast. For chromatic directions that had little-to-no L cone contrast, the lag versus contrast functions were decreasing at the highest contrasts available within our display gamut. We quantify the relative contribution of the L- and S-cones cones to tracking performance in the section below.

### Experiment 1: Color Tracking Model (CTM)

We developed a color tracking model (CTM) that relates the L- and S-cone contrast of a stimulus modulation to tracking lag. The first stage of the model combines contrast from the two cone mechanisms through a quadratic computation. This stage transforms stimulus contrast and chromatic direction into what we call equivalent contrast. This is the effective contrast of the stimulus after it has been weighted by the sensitivity of the underlying mechanisms for the corresponding chromatic direction. The first stage can be summarized by an elliptical contour whose shape indicates the relative L- and S-cone contrasts in any chromatic direction that lead to the same equivalent contrast. In the CTM, the shape of the elliptical isoresponse contour is constrained to be independent of overall contrast.

This equivalent contrast computed by the first stage of the model provides a common axis for the lag measurements, collapsed across chromatic direction. The second stage of the model uses a single exponential decay function to transform equivalent contrast to tracking lag. This function is independent of the chromatic direction of the stimulus, as it depends only on the equivalent contrast. Hence, the elliptical contour that characterizes how equivalent contrast depends on chromatic direction specifies an isoresponse contour. It describes the relative L- and S-cone contrasts in any chromatic direction that lead to the same tracking lag.

The shape of isoresponse contours is specified by two parameters: 1) the ellipse angle *θ* (representing the direction of least sensitivity; counterclockwise to the positive abscissa), and 2) the minor axis ratio *m* (the ratio of vector lengths between the most and least sensitive directions). Within this framework, these parameters provide a full account of the dependence of performance on chromatic direction for the tracking task. Figure 3 shows the isoresponse contours for the CTM for all three subjects, with the parameter values for each subject inset in each panel. These are shown with their major axes normalized to have a length of 2 (±1 in each direction around the origin).

**Figure 3:**
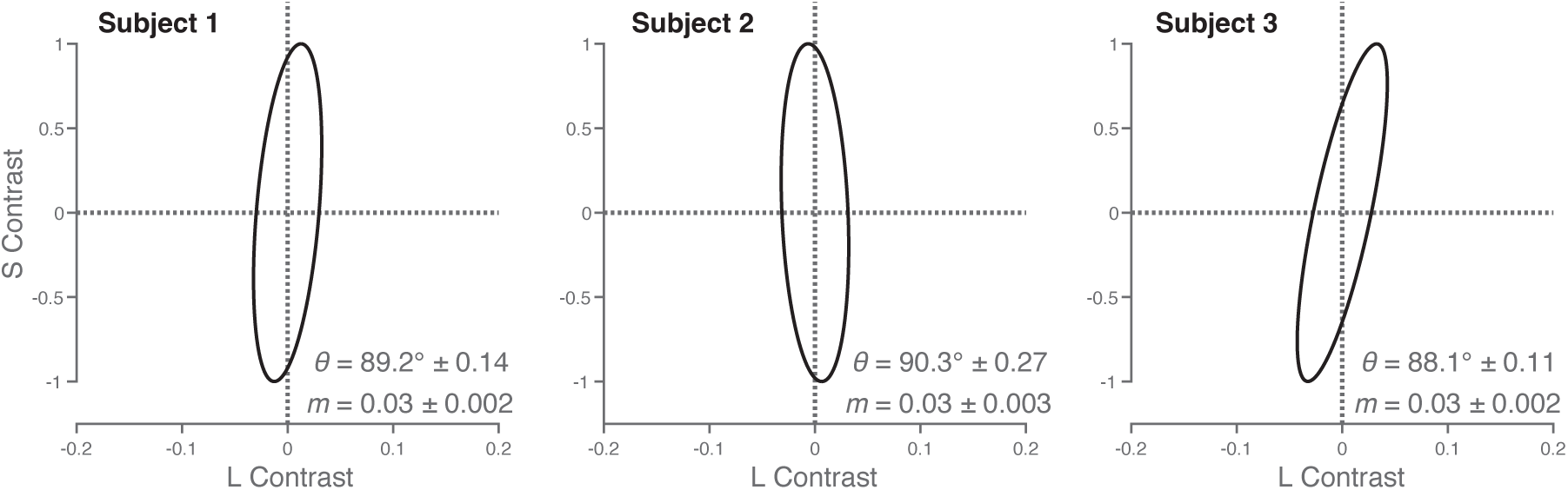
Isoresponse Contours for the Color Tracking Model. The grey ellipse in each panel shows the isoreponse contour associated with the color tracking task for each subject. The isoresponse contour is the set of stimuli that result in the same tracking lags. This contour is constrained in the CTM to take the form of an ellipse in the LS cone contrast plane, and its shape is constrained to be the same independent of overall contrast. The ellipse is specified by two parameters: 1) the ellipse angle and 2) the minor axis ratio. The ellipse angle represents the direction of least sensitivity defined counterclockwise to the positive abscissa. The minor axis ratio is the ratio of the vector lengths between the most and least sensitive directions. The scale of the ellipses across subjects is normalized to have a length of 2 along their major axis (1 in each direction). Also note the expanded scale of the L-contrast axis. The ellipse parameters (means and their standard errors estimated by bootstrap resampling) are provided in the insets.

For all subjects, *θ* was close to 90°, implying that the chromatic mechanisms underlying tracking are least sensitive to stimuli modulated in the S-cone direction (and most sensitive to stimuli modulated in the L-cone direction). The parameter *m* captures the magnitude of this difference in sensitivity across chromatic directions. All three subjects had an *m* value of 0.03, implying that tracking performance as summarized by tracking lag is ∼30x more sensitive to L-cone isolating stimuli than to S-cone isolating stimuli.

Because the empirically determined ellipses are quite distended, we took special care to choose stimuli aligned with the least sensitive direction (see Horiguchi, Winawer, Dougherty, & Wandell, 2013 for a similar approach). After an initial determination of the ellipse shape from measurements made with a common set of stimuli for each subject, we tailored additional stimuli to preliminary model fits for each subject, so that estimates of ellipse size and orientation would be well constrained by the data.

The isoresponse contour can be used convert stimuli of varying contrast and chromatic directions into equivalent contrast. This value for each stimulus was then related to tracking lag via a single non-linear function (a three-parameter exponential decay) that was shared across all chromatic directions. Figure 4 presents the form of the response nonlinearity that was found for each subject. In all subjects, an increase in equivalent contrast was associated with a decrease in lag, reaching an asymptote of approximately 350 ms in all three subjects. The best-fit parameter values (*A*, *s*, and *d*; see Methods for functional form) differed between subjects and are provided in each panel of Figure 4. The insets indicate the subject specific chromatic directions used to help constrain the ellipse parameters for each subject.

**Figure 4:**
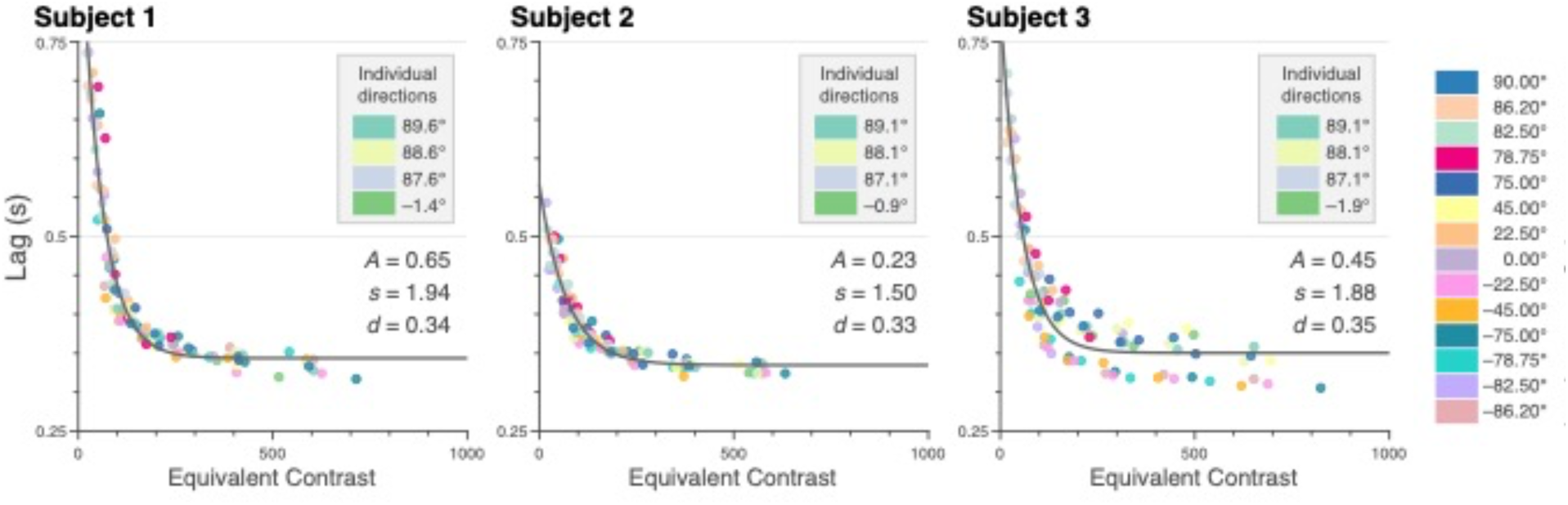
Nonlinearity of the Color Tracking Model. The grey curve in each panel shows the nonlinearity of the color tracking model. The x-axis it the equivalent contrast which is the result of the isoresponse contour. The y-axis respresent the response, which in this figure, is the lag from the tracking task in seconds. The closed circles in each plot are the tracking lags from the tIRF. The stimulus contrasts of the for each lag has been adjusted by the isoresponse contour allowing the lags for all stimuli to be plotted on the equivalent contrast axis. The color map denotes which color correspond to which directions. The inset color map in each panel marks the directions tested which were unique to each subject.

The ability of the CTM to account for the lag data is summarized by the agreement between transformed data and the response nonlinearity (Figure 4) as well as by the dashed fit lines to the untransformed tracking lags (Figures 2 and S1). Indeed, the five-parameter model (two for the isoresponse contour; three for the non-linearity) provides a good account of tracking performance across contrast levels and chromatic directions. One exception was found in Subject 3, for whom there appeared to be some bifurcation of transformed lag values relative to prediction at high equivalent contrast levels (Figure 4, left panel). We do not have an explanation for this.

### Experiment 2: Chromatic contrast and detection

Subjects participated in a two-interval forced choice (2IFC) task in which they were asked to report which of two intervals contained a moving Gabor target. While it was presented, this target underwent random walk motion with the same parameters used in the tracking experiment. The Gabor targets varied in chromatic direction and contrast as in the tracking task, although we used fewer chromatic directions in this experiment. We measured fraction correct detection for each direction and contrast. As with to the tracking lag data, we examined the relationship between fraction correct and contrast by grouping the measurements by their chromatic direction. For all subjects and chromatic directions (Figures 5 and S2), fraction correct increased with stimulus contrast.

**Figure 5:**
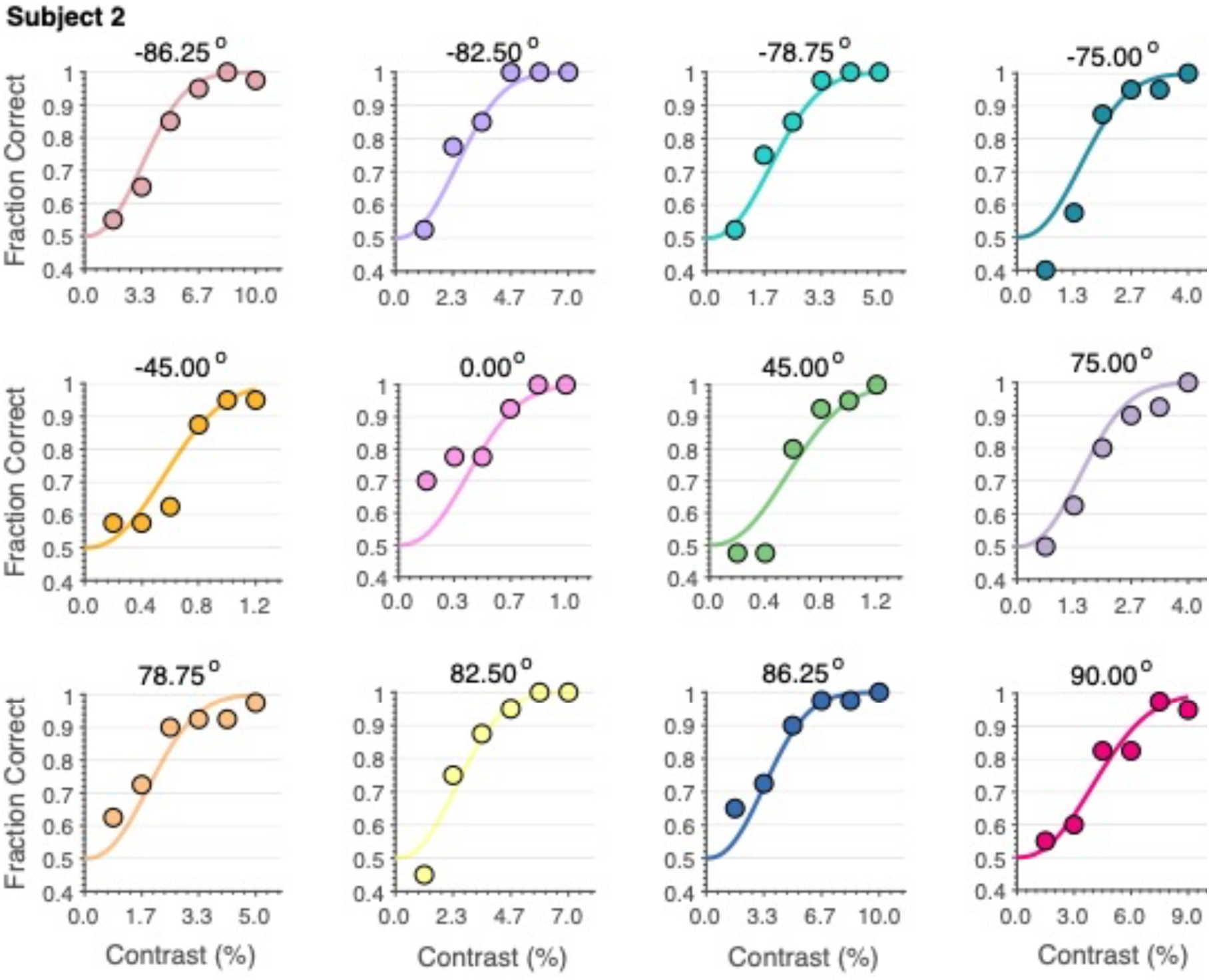
Detection Versus Contrast for Subject 2. The figure shows the fraction correct in the detection task as a function of the stimulus contrast for each chromatic direction used in the experiment. The closed circles in each panel are the fraction correct for an individual stimulus direction, with angle in the LS contrast plane provided above each panel. The solid curves are the fraction correct predictions of the color detection model fit to all the data simultaneously (see below). Note the difference in x-axis range between panels.

Differences in sensitivity across the chromatic directions may be appreciated by considering the stimulus contrast required to achieve a criterion detection threshold (e.g., 76% correct). For L-cone isolating stimuli, this contrast is less than a percent, while it is several percent for S-cone isolating stimuli. The very high sensitivity to L-cone isolating modulation suggests that this stimulus is detected by an L-M cone-opponent mechanism (Chaparro, Stromeyer, Huang, Kronauer, & Eskew, 1993).

### Experiment 2: Color Detection Model (CDM)

We developed a model of detection performance as a function of stimulus direction and contrast in the LS plane. As with the CTM, the CDM is also composed of two stages, with the first based on a quadratic computation of equivalent contrast (elliptical isoresponse contours), which in this case identifies the stimulus contrast in each chromatic direction that produces the same level of detection performance. The second stage is a cumulative Weibull (specified for a guessing fraction correct of 0.5). This allowed us to convert the equivalent contrast to a prediction of fraction correct, bounded between 0.5 and 1.0.

The elliptical isoresponse contours of the CDM for each subject are plotted in the panels of Figure 6. The model and plot conventions are the same as used in Figure 3. For all subjects, *θ* was again close to 90°. Although the differences in angle across the two experiments exceed the measurement confidence as assessed by bootstrapping, the magnitude of the numerical differences (less than 2 degrees) are sufficiently small in an absolute sense that we do not attribute meaning to these differences.

**Figure 6:**
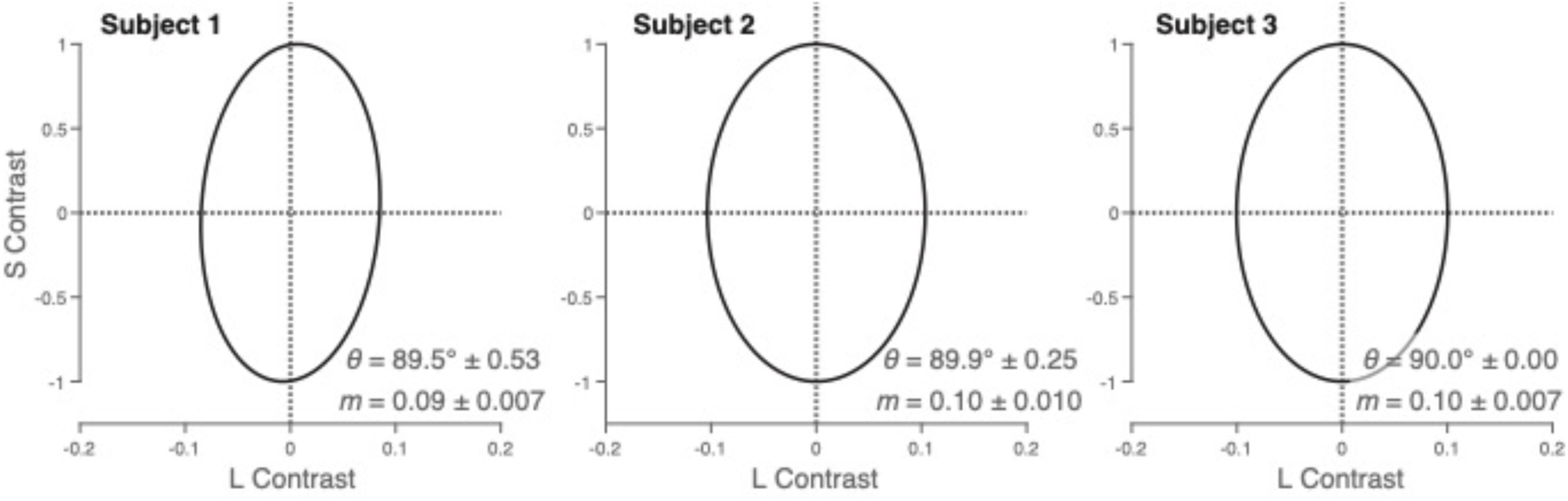
Isoresponse Contours for the Color Detection Task. The grey ellipse in each panel shows the isoreponse contour associated with the color detection task for each subject. These contours define a set of stimuli which produce the same fraction correct in the detection task. The isoresponse contour of the color detection model has the same parameterization as the color tracking model. The scale of the ellipses across subjects is normalized to have a length of 2 along their major axis (1 in each direction). Also note the expanded scale of the L-contrast axis. The ellipse parameters (means and standard errors estimate by bootstrap resampling) are provided in the insets.

The minor axis ratios (*m*) for the threshold experiment ranged between 0.09 and 0.10 across the subjects, implying that the underlying chromatic mechanisms for detection were ∼10x more sensitive to L-cone isolating stimuli than S-cone isolating stimuli. Recall that, for tracking performance, relative sensitivity to L-cone isolating stimuli was considerably stronger (∼30x rather than ∼10x). This difference in relative sensitivity is substantial, and well beyond the uncertainty in parameter estimation as established by bootstrapping.

Figure 7 shows the nonlinearity of the CDM that relates equivalent contrast to detection performance. An increase in equivalent contrast was associated with an increase in detection performance that was well described by the function. There were slight variations in the values of the two parameters that define the function for each subject (shown inset in each panel).

**Figure 7:**
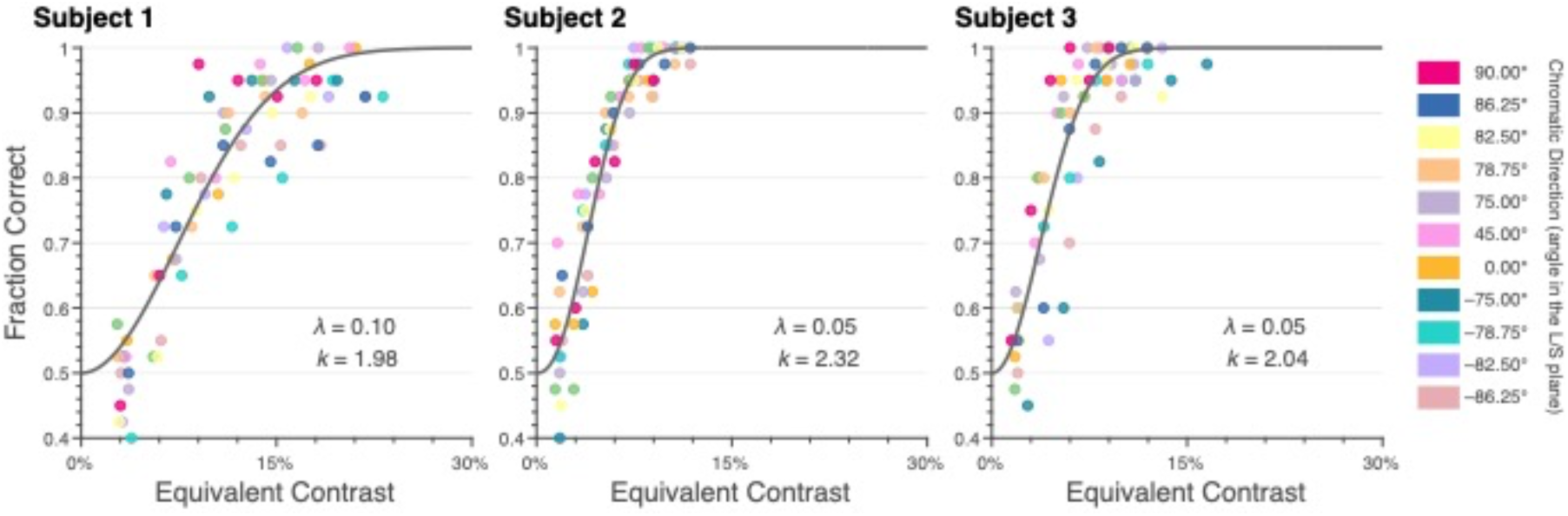
Nonlinearity of the Color Detection Model. The grey curve in each panel shows the nonlinearity of the color detection model for all subjects. The x-axis is equivalent contrast. The y-axis is the response, which in the figure, is the fraction correct of the detection task. The closed circles in each plot are the fraction correct for each condition tested. The stimulus contrast for each closed circle has been adjusted by the isoresponse contour, allowing the fraction correct to be plotted on the equivalent contrast axis. The color map denotes which color corresponds to which direction.

The ability of the CDM to account for the detection data is demonstrated by the overall fit to the data shown in Figure 7 as well as solid fit lines shown in Figures 5 and S2. The four-parameter model (two for the isoresponse contour; two for the non-linearity that transforms equivalent contrast to fraction correct) provided a good account of detection performance across contrast levels and chromatic directions.

To illustrate how the CTM may be used to re-express the results in interesting ways, we computed the predicted difference in tracking lag between S- and L-cone isolating stimuli, when the cone contrast across the two directions was equated. Figure 8 shows the result of this calculation for each of the three subjects. Each subject shows the same qualitative dependence: in each case the lag difference decreases systematically with contrast. The fact that lag difference depends strongly on cone contrast emphasizes the importance of jointly characterizing the dependence of performance on both chromatic direction and contrast. More generally, the contrast dependence of the relative lag means reminds us that it important to consider performance at multiple contrast levels: the effects of an independent variable (here color direction) can depend considerably on contrast.

**Figure 8:**
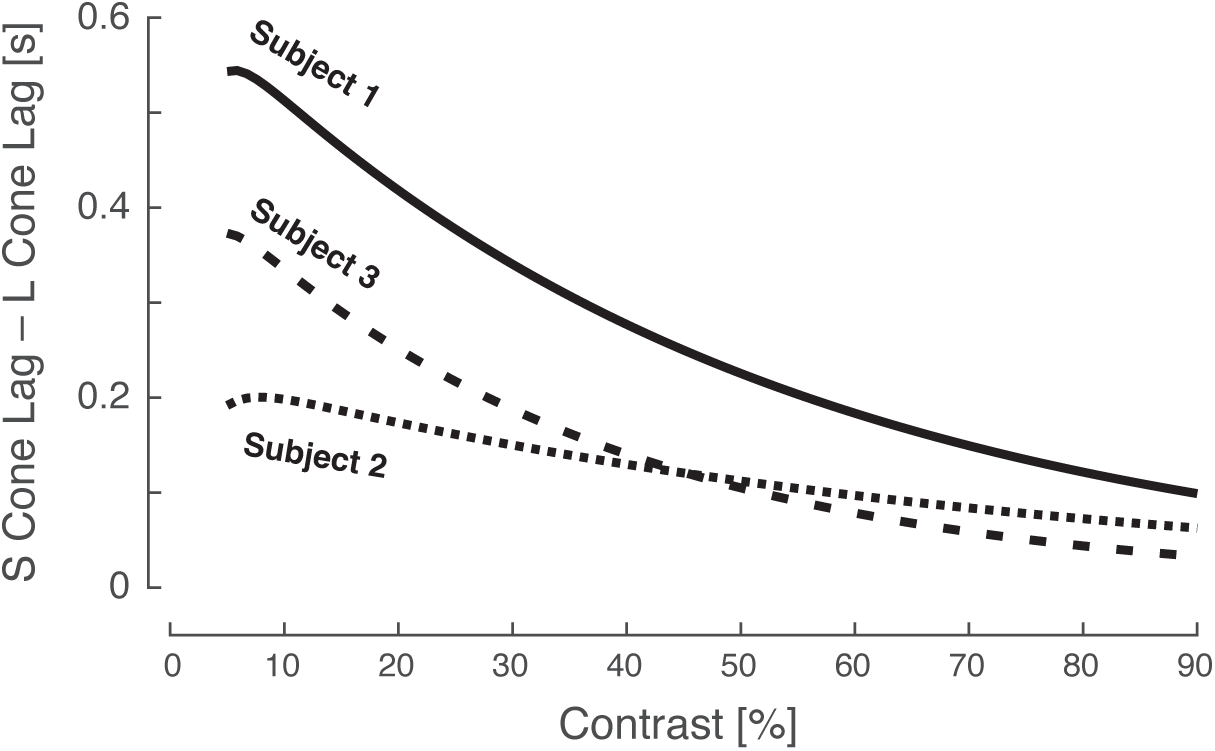
Comparison of predicted S- and L-cone tracking lags. The figure shows predictions obtained from the CTM model for the difference between S- and L-cone tracking lag, when contrast is equated across the two chromatic directions. The x-axis shows the common cone contrast, and the y-axis the difference in tracking lag in seconds. The model would allow similar predictions to be made for any pair of chromatic directions and relative contrasts.

## Discussion

We measured tracking and detection performance for a set of matched stimuli that varied in chromatic direction and contrast. We find that each task can be accounted for by a two-stage model. The two models, the CTM and the CDM, each have first stages of the same form: a quadratic combination of cone contrast that computes an overall equivalent contrast. The second stage is a task-specific, non-linear readout that relates equivalent contrast to performance. This model structure, which separates effects of chromatic direction from those of contrast, characterizes the role of chromatic direction on performance in a contrast-independent manner. Such separability between chromatic direction and contrast is a prerequisite for making well-defined statements about the sensitivity of performance as a function of the chromatic direction of the modulation. If performance were not direction-contrast separable, then consideration of the effect of chromatic direction would need to be accompanied by specification of what contrasts were being considered for each direction. The elliptical contours provided in Figures 3 and 6 summarize our findings about how performance depends on chromatic direction for each task.

Similarly, the direction-contrast separability also allows us to make statements about the dependence of performance on contrast that are independent of chromatic direction. Figures 4 and 7 summarize our findings about this dependence for each task.

Quadratic models have been used previously to understand color sensitivity (e.g., Guth, Massof, & Benzschawel, 1980; Krauskopf, Williams, & Heeley, 1982; Poirson, Wandell, Varner, & Brainard, 1990; Poirson & Wandell, 1990b; Poirson & Wandell, 1990a; Guth, 1991; Poirson & Wandell, 1993; Poirson & Wandell, 1996; Knoblauch & Maloney, 1996), color motion processing (Chichilnisky, Heeger, & Wandell, 1993), and cortical responses to colored stimulus modulations (Horwitz & Hass, 2012; Barnett, Aguirre, & Brainard, 2021).

If the iso-performance ellipses for the two tasks had had the same shape, we could have concluded that the same combination of cone signals (i.e. the same equivalent contrast) accounted for performance in both tasks. This result was not observed. Instead, there was a substantial difference in relative sensitivity for S as compared to L chromatic directions, with a deficit for S-cone isolating stimuli of ∼3x observed for tracking relative to detection.

Although the iso-performance contours differed in aspect ratio in the two tasks, the orientation of the iso-performance contours in the two tasks was essentially the same. This is consistent with the notion that performance on the two tasks is mediated by the same mechanisms, but with the sensitivity of the readout of S cones reduced for tracking relative to detection. Because of the design of our stimuli, which matched temporal properties across the two tasks, the difference in sensitivity is not due to differences in the temporal properties of the moving stimuli *per se*.

Although our experiments do not uniquely determine the underlying mechanisms,^1^ it seems likely that detection of L-cone isolating stimuli is mediated by the highly-sensitive L-M cone opponent mechanism, while detection of S-cone isolating stimuli is mediated by the S-(L+M) mechanism. Tracking for our stimuli could then be mediated by the same two mechanisms. Our data do not, however, conclusively rule out the possibility that the mechanisms that supporting tracking differ from those that support detection. As one example, tracking L-cone isolating stimuli could mediated by the L+M cone mechanism rather than the L-M cone mechanism, or a combination of the two. Experiments that systematically modulate M- as well as L-cone contrast would help resolve this ambiguity.

Under the reasonable assumption that tracking performance is related to the ability of a subject to perceive stimulus motion, then the S-cone deficit in tracking is consistent with previously observed motion processing deficits for S-cone isolating stimuli (Cavanagh, MacLeod, & Anstis, 1987; Dougherty, Press, & Wandell, 1999). Similarly, a delay in signals originating with the S-cones as revealed by earlier studies (Smithson & Mollon, 2004; McKeefry, Parry, & Murray, 2003) would also be expected to produce an S-cone tracking deficit. In addition, we emphasize that our approach provides an account which generalizes across chromatic directions (here confined to the LS contrast plane) and across contrasts.

Future work would benefit from extending measurements beyond a single plane in color space to characterize the full, three-dimensional iso-response contour. Such contours would be expected to be well-described by ellipsoidal quadratic forms (Poirson, Wandell, Varner, & Brainard, 1990; Knoblauch & Maloney, 1996). Indeed, if measured in a full set of directions in three-dimensional LMS contrast space, the logic introduced by Poirson and Wandell (Poirson & Wandell, 1993; Poirson & Wandell, 1996) could be used to leverage the change in sensitivity across tasks to estimate the inputs to a set of task-mediating cone-opponent mechanisms. Similarly, changes in the spatial frequency and motion parameters of the tracked stimuli could be exploited to further generalize the findings and constrain inferences about underlying mechanisms.

The current work constitutes an important step towards a broad goal of predicting how the temporal dynamics of visual processing and visuomotor behavior (e.g. the tIRFs) depend on the spatio-chromatic properties of stimuli. Future experiments should aim to understand how the particular contrasts, colors, and spatial frequencies defining an arbitrary stimulus determine these dynamics.

## Materials and Methods

### Subjects

Three subjects (ages 28, 29, 33; two male) took part in all psychophysical experiments. All subjects had normal or corrected to normal acuity and normal color vision. All subjects gave informed written consent. The research was approved by the University of Pennsylvania Institutional Review Board. Two of the subjects are authors on this paper; one was naïve to the purpose of the study.

### Experimental Session Overview

All subjects participated in both the color tracking task and color detection task experiments. In total, all experiments spanned eight sessions. The first six sessions were for the tracking task, and the remaining sessions were used for the detection task. Each of the tracking sessions lasted approximately 1.5 hours and each detection session lasted approximately 1 hour. Subjects completed the full set of tracking sessions before starting the detection sessions. All experiments were preregistered: the Color Tracking Task (Exp. 1: https://osf.io/xvsm3/; Exp. 2: https://osf.io/5y2dh/; Exp. 3: https://osf.io/e6dfs/) and the Color Detection Task (Exp. 4: https://osf.io/ekv24/; Exp. 5: https://osf.io/ekv24/).

### Stimulus Display and Generation

The stimuli were designed to create specific responses of the cone photoreceptors using silent substitution (Estévez & Spekreijse, 1982). Silent substitution is based on the principle that sets of light spectra exist that, when exchanged, selectively modulate the activity of cone photoreceptors. Therefore, modulating the stimuli, relative to a background, can selectively modulate the activity of the L-, M-, or S-cones, or combinations of cones for a specified contrast. These calculations require a model of the spectral sensitivities of the cone photoreceptors and spectral power distributions of the monitor primaries. We use the Stockman-Sharpe 2-degree cone fundamentals as the model of the cone sensitivities. To obtain the spectral power distribution of the monitor RGB primaries, we performed a monitor calibration using a PR-650 SpectraScan radiometer. This also allowed us to obtain the gamma function for each primary as well.

All stimuli were generated using a ViewSonic G220fb CRT monitor with three primaries and had a refresh rate of 60 Hz. The horizontal and vertical resolution of the monitor was 1024x768 pixels, respectively, corresponding to a screen size of 405 x 303 mm. Subjects viewed the monitor at distance of 92.5 mm for tracking and 105 mm for detection.

### The Visual Stimuli

The stimuli for all experiments were restricted to the LS plane of cone contrast space. Cone contrast space is a three-dimensional space with each axis showing the change in the quantal catch of the L, M, and S cones relative to a specified reference spectrum. This reference spectrum is referred to as the background light. The background used in all experiments was luminance: Y = 30.75 cd/m2, chromaticity: x = 0.326, y = 0.372. These were calculated using the XYZ 2° color matching functions, https://cvrl.org). We set the origin of the LS cone contrast plane to be this background and confined all modulation to this plane. Modulations made around this background have the effect that they only modify the L- and S-cone excitations, while leaving M-cone excitations unchanged. We refer to the chromatic component of the stimuli used in this experiment as vectors in this plane. Each stimulus has an L-cone and S-cone vector component, and we refer to the stimuli by the angle computed by the ratios of these components. In this space, S-cone isolating stimulus vectors are oriented at 90° and L-cone isolating stimulus vectors are oriented at 0°. We refer to these angles as the chromatic directions. The contrast of a stimulus is defined as the L2-norm of the stimulus vector in the LS plane.

The spatio-temporal parametrization of the stimuli are identical across both experiments. The stimuli used were sine phase Gabor patches. The frequency of the sine wave was set to 1 C.P.D. and the standard deviation of the Gaussian window was set to 0.6 degrees of visual angle (DVA). This standard deviation corresponds to a FWHM of 1.41° and 90% of the Gaussian envelope being contained in a 2° diameter window. The Gabor patch performed a random walk confined to move horizontally across the monitor. The Gabor target updated its position on each frame according to a Gaussian velocity distribution. This distribution was centered on 0 °/s with a standard deviation of 2.6 °/s and a FWHM of 6.1 °/s. Values drawn from this distribution with a negative sign corresponded to leftward motion and values with a positive sign corresponded to rightward motion. This resulted in an average speed of 2.07 °/s and an average step size of approximately 0.63 mm. What varied across stimuli was the chromatic content of the Gabor with the spatio-temporal parameters fixed for all directions and contrasts. The exact chromatic directions and contrasts used in each experiment are reported their respective sections below.

### The Color Tracking Task

Subjects participated in the continuous tracking task in which they were asked to track the position of a target Gabor patch. On each trial, the Gabor patch spatially jittered its position along a horizontal linear path across the middle of the monitor in accordance with the temporal parameters noted in the prior section. The subjects were instructed to indicate the position of the Gabor patch by continuously trying to keep the cursor in the middle of the patch. Subjects controlled the position of the cursor though the use of a computer mouse. At the end of each trial, we obtain a time-course of the target positions on the screen and the subjects cursor responses. Each trial lasted 11 seconds with an initial static period of one second. Example traces of the target (grey line) and tracking (black line) position as a function of time can be seen in the upper portion of panel c of Figure 1.

The Gabor patches were modulated in 18 different chromatic directions each with 6 contrast levels. Tables S1 and S2 summarize the directions and contrasts used in the Color Tracking Task for all subjects. These directions were split into three sets of experiments each with 6 directions. Within each set, subjects completed 2 sessions each made up of 10 experimental runs. An experimental run consisted of 36 trials corresponding to a single presentation of each of the conditions (6 directions and 6 contrast). Across the runs, we flip the handedness of the sine-phase Gabors such that alternating runs are offset by 180° spatial phase of the Gabor. The order of trials within a run was pseudo-randomized such that each session contained the desired number of repeats. Subjects controlled the pace of the trials and between trials only the background was present. A single session contained 360 trials lasting approximately 1 hour. In total, across all three experimental sets, 2,160 trials were collected per subject equal to 6 hours of tracking data.

### The Temporal Impulse Response Function

From the time-courses of the target and the tracking positions, we can compute the temporal impulse response function associated with the color tracking task for each chromatic direction and contrast condition within an individual subject. In a linear system, the temporal impulse response function is the response of the system, as a function of time, to a brief presentation of the input. In the context of the color tracking task, the tIRF describes the response of the visual-motor system to the movements of the random walk of the Gabor target. To obtain the tIRF, we perform a cross-correlation between the velocities of the target’s random walk and the velocities of the subject’s tracking. The velocities of the targets have a white noise power spectra containing no temporal autocorrelations. Example velocity traces for a given run can be seen in the lower portion of panel c of Figure 1. In this panel, the target velocity is shown as the grey line and the tracking velocity is shown as the black line.

If the subject were to perfectly track the Brownian motion of the target, we would end up with two identical white noise velocity traces. The cross correlation of two white noise signals produces a delta function centered on 0 meaning they are perfectly correlated only when the signals are temporally aligned and have no other correlation as a function of delaying one signal relative to the other. Now if the subject perfectly tracked the target motion but had a consistent 2 second delay in their tracking then the resulting cross-correlation function would again be a delta function but centered at 2 second rather than 0. Deviation from the delta function shape is due to factors such as noisy tracking or dependance on the stimulus history to inform of the target’s current position. In the current experiment, we use the tIRFs to inform us of the delay of the visual-motor system associated tracking.

The tIRFs are obtained via the cross-correlation of the concatenation of position and tracking for all runs with a stimulus condition. This provides us with a mean tIRF for each chromatic direction and contrast pair. Example mean tIRFs are plotted as the grey lines in panel d of Figure 1. This shows the correlation between the two signals as function of delaying one relative to the other for a particular direction and contrast. To reliably estimate lag from the mean tIRF, we fit a log-Gaussian function to the cross-correlogram and take its mode as the lag of the tIRF. This tells us the time at which the log-Gaussian reaches its maximum correlation. An example log-Gaussian fit can be seen as the purple curve in panel d of Figure 1. Overall, we obtain one lag estimate per stimulus condition. There is additional information in the tIRF beyond lag, that we did not examine here.

The error bars on the lag estimates plotted in Figure 2 are computed as the standard deviation of bootstrapped lag estimates, which provides an estimate of the SEM of these estimates. This same bootstrapping approach was used to estimate SEMs in the other cases where measurement precision is reported.

### The Color Tracking Model

The Color Tracking Model (CTM) is a 5-parameter model that provides a prediction of tracking lag for any input stimulus specified in terms of its L- and S- cones contrasts. A detailed formulation of a related model maybe found in the model appendix of Barnett et al. 2021. Converting chromatic direction and contrast into lag is done through two stages. The first stage is the application of a quadratic isoresponse contour. The isoresponse contour effectively weights the input stimulus contrast by the underlying chromatic mechanism sensitivity for the corresponding chromatic direction.

This weighting produces a single output variable from the L- and S-cone contrast inputs, collapsing across chromatic direction. We refer to this output variable as ‘equivalent contrast’ and it represent the strength of the chromatic mechanism output which is now independent of the original chromatic direction. The isoresponse contour represents sets of stimuli that when shown to a subject result in equal tracking lags. In the CTM, the shape of the isoresponse contour is constrained to be an ellipse restricted to the LS cone contrast plane.

Of the 5 parameters in the CTM, 2 of them are used to specify the elliptical isoresponse contour. One of these parameters is the ellipse angle. The ellipse angle represents the direction of least sensitivity in the LS plane and read counterclockwise to the positive abscissa. Since the parametrization of the model enforces two orthogonal mechanisms, the second mechanism represents the direction of maximal sensitivity.

The other parameter is the minor axis ratio. This is the ratio of the minor to major axis vector lengths. In this model, the length of the major axis is locked to unit length and the minor axis is constrained to be less than the major axis. Within this, minor axis vector length can be readily interpreted as this ratio. Together, these parameters provide a complete account of the chromatic stage of the model.

The second stage of the CTM employs a single nonlinearity which transforms the equivalent contrast to a prediction of the tracking lag. Since the isoresponse contour allows us to collapse across chromatic directions to a scalar equivalent contrast, we can use a single nonlinear function to map to tracking lag. The functional form we employ to convert equivalent contrast to lag is a the three-parameter exponential decay function:

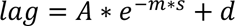

In this expression, the parameters are *A*, *s*, and *d* which represent the amplitude, scale, and offset, respectively. The scale parameter operates on the equivalent contrast (*m*) and acts as gain on the output of the chromatic stage of the model. The offset (*d*) is interpreted as the minimum lag, this is the point at which the decay function asymptotes.

### The Color Detection Task

Subjects participated in a color detection task which allowed for estimates of detection thresholds to be obtained for each chromatic direction. The detection task used a two-interval forced choice task paradigm in which a Gabor stimulus was presented in one of two sequential intervals. The start of each interval was marked with a brief tone as well as the disappearance of the fixation dot. Each interval had a duration of 400 ms and were separated by a 200 ms gap between intervals in which the fixation dot was present. At the end of the trial, the subject indicates via a gamepad button press the interval in which they think the stimulus was presented. Based on this response, the subject receives auditory feedback (high pitch tone for correct, low pitch tone for incorrect).

The Gabor stimuli in the detection task had the same spatio-temporal properties as those used in the tracking task. Therefore, the stimuli, during their presentation, performed the same random walk for 400 ms. The primary way in which the non-chromatic properties of the stimuli differed across tasks was in the stimulus ramping. At the beginning and end of the interval containing the target, the contrast of the Gabor was temporally windowed. This window was a half-cosine ramp with a duration of 100 ms. The structure of the target interval was as follows: a 100 ms ascending ramp at the beginning, 200 ms of full stimulus contrast, and a 100 ms descending ramp at the end.

For this experiment, we modulated the Gabor stimuli in 12 chromatic directions, these directions being a subset of the 18 directions used in the tracking. These chromatic directions were modulated around the same background across the two tasks. Within each chromatic direction, we tested at 6 evenly spaced contrast levels between the maximum contrast and 0 (excluding 0). The maximum contrasts were determined in pilot experiments for each subject and are intended to effectively sample the rising portion of the psychometric function. The directions tested and their corresponding maximum contrasts are reported in Table S3.

The stimuli were displayed on the same CRT monitor as the previous experiments. One difference is the need for finer control of contrast than was needed for the previous tracking experiments. To achieve the required bit depth, we used a Bits++ device (Cambridge Research Systems) to enable 14-bit control of the R, G, and B channel inputs to the CRT. In addition, a different host computer was used to control the experiment (Asus RoG laptop and Ubuntu 20.04) for compatibility with the Bits++ device.

Trials were blocked so that 60 trials of a given direction will be shown consecutively. Each block contained 10 presentations of each of the contrast levels. The contrasts are pseudorandomized such that a random permutation of all 6 levels were shown before repeating a contrast. To orient the subject to the chromatic direction of the block, there were 3 practice trials shown at the start of each block for the highest contrast level. Subjects completed a total of 40 trials per contrast/direction pair, that is 4 blocks per direction. Half of the blocks for each direction will be left-handed Gabors and the other half will be right-handed Gabors. We have 12 directions each with 6 contrast levels for a total of 240 trials per direction and a total of 2,880 trials.

### Threshold Detection

From the detection data, we estimate a threshold for each of the chromatic directions tested. Threshold is the stimulus contrast needed, per direction, to reliably detect the Gabor target. We estimate this value by fitting a psychometric function to the fraction correct as a function of stimulus contrast for each direction. Specifically, we fit a cumulative Weibull function and use it to determine the contrast needed to reach 76% correct.

### The Color Detection Model

The Color Detection Model (CDM) is a 4-parameter model that provides a prediction of the fraction correct in the detection task for any input stimuli specified in terms of its L- and S- cones contrasts. The CDM parallels the CTM in its construction. It also employs two stages in order to convert stimulus contrast to fraction correct. The first stage is an elliptical isoresponse contour identical to the one used in the CTM (see The Color Tracking Model section). Since the relationship between equivalent contrast and the variable of interest across the two tasks have different forms, we need to employ task-dependent nonlinearities as the second stage. In the CTM, this relationship was captured with an exponential decay function. For the detection task we use the cumulative Weibull as the nonlinearity that converts the equivalent contrast into fraction correct. The functional form we use is:

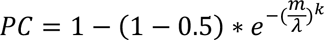

In this expression, the parameters are 𝜆 and *k* which represent the scale, and shape, respectively. The scale parameter operates on the equivalent contrast (*m*) and acts as gain on the output of the chromatic stage of the model. The shape parameter (*k*) controls the slope. The guess rate is locked at 0.5 since this is chance in a 2IFC, therefore the part of the expression ‘(1-0.5)’ bounds the output fraction correct between 0.5 and 1.

### Parameter Fitting

We fit both the CTM and CDM to their respective measurements as a function of their stimulus contrasts. These data were fit using the MATLAB function *fmincon* to find a set of model parameters that minimize the root mean squared error between the actual tracking lags or fraction correct and the predicted values of the CTM and the CDM, respectively.

---

**Figure S1:**
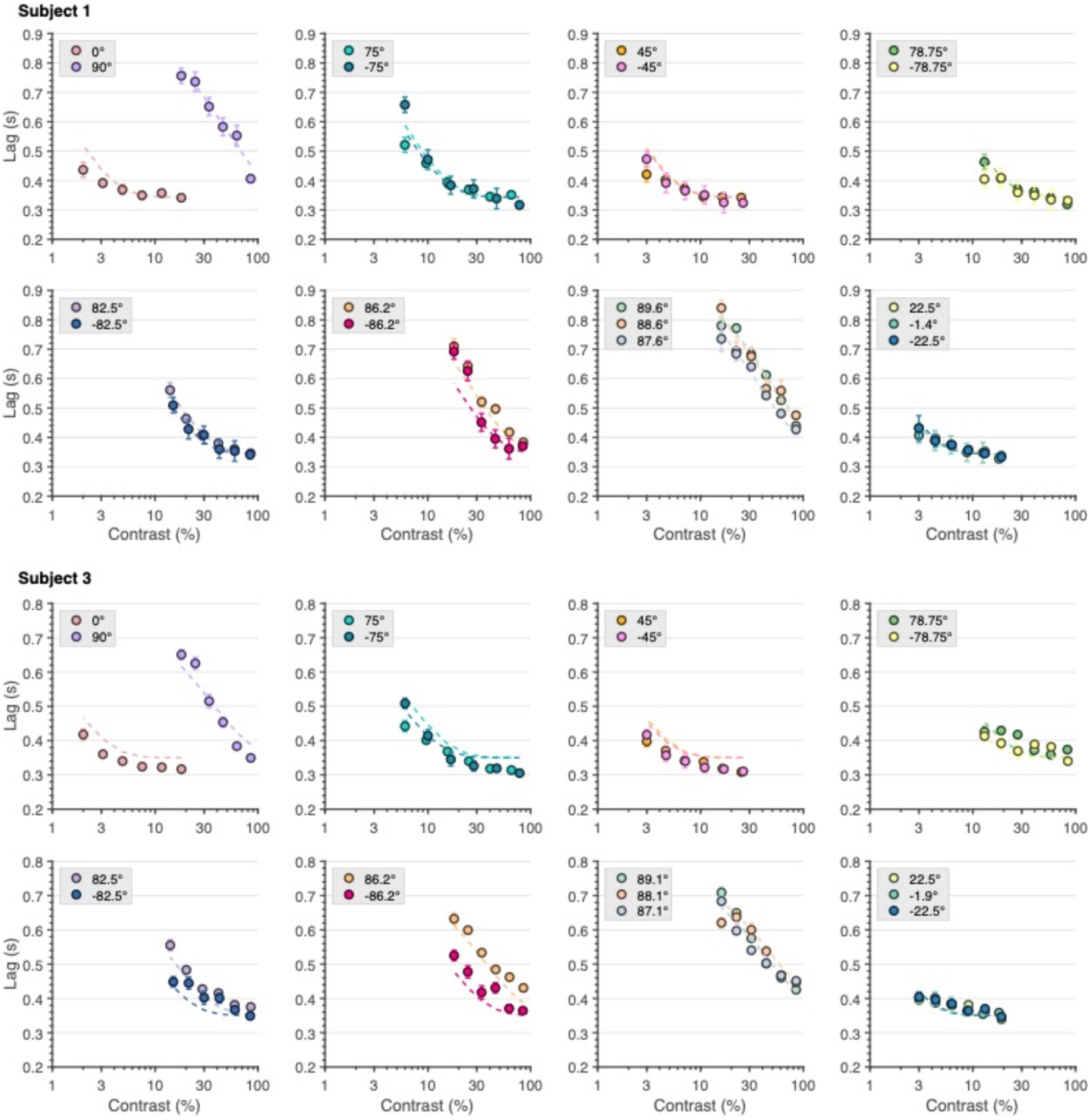
Tracking Lag Versus Contrast for Subjects 1 and 3. Same format as Figure 2.

**Figure S2:**
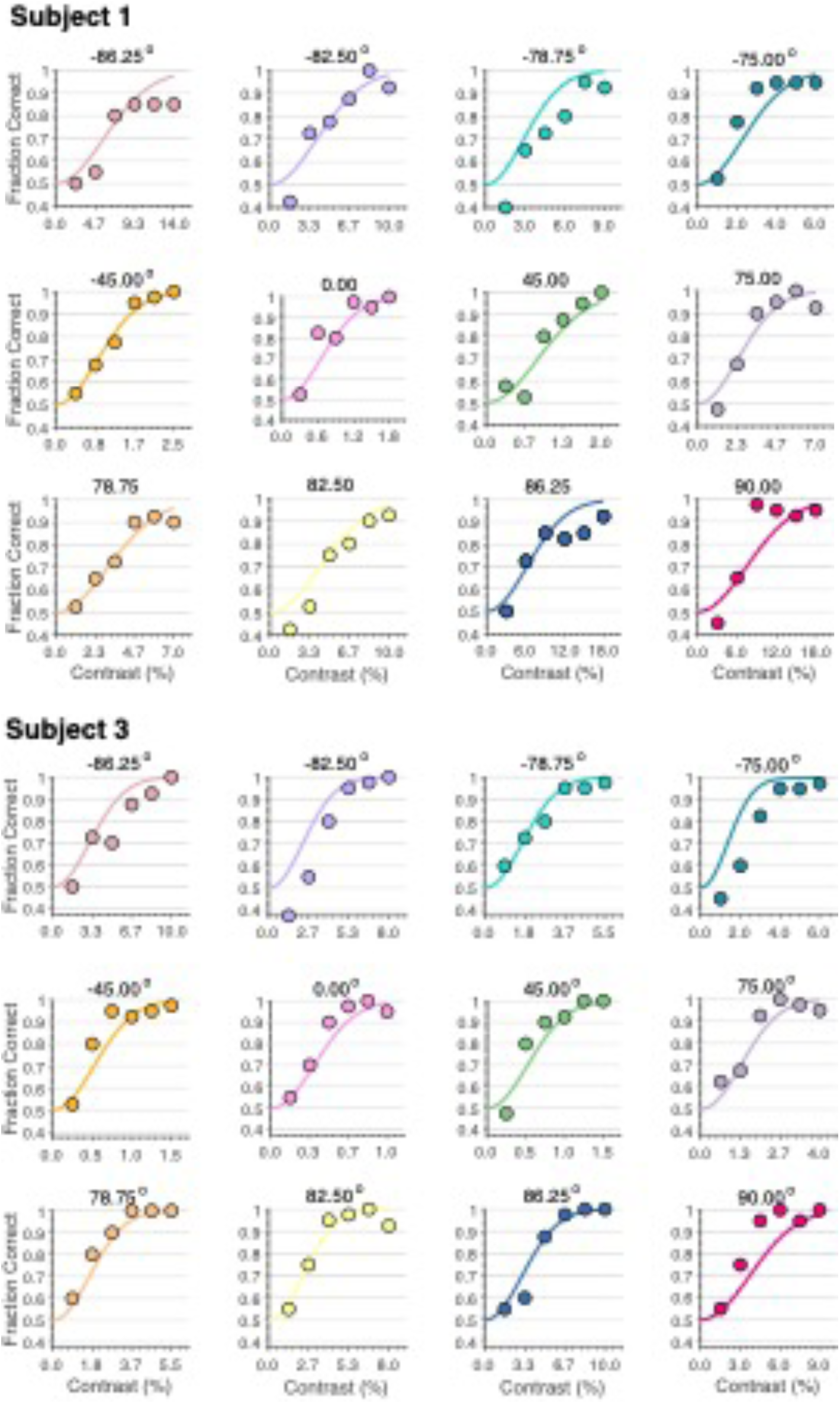
Detection Performance Versus Contrast for Subjects 1 and 3. Same format as Figure 5.

**Table S1:**
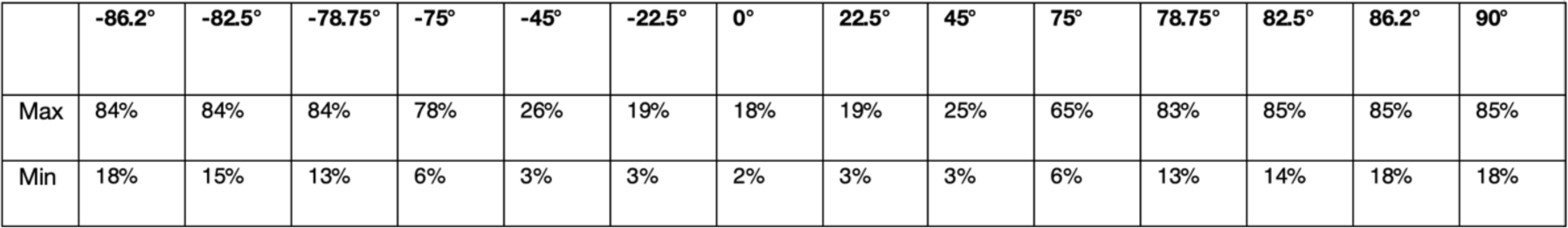
The Chromatic Directions and Contrasts – CTM. The chromatic directions and maximum/minimum contrasts of the tracking task. This table includes the chromatic directions common to all subjects.

**Table S2:**
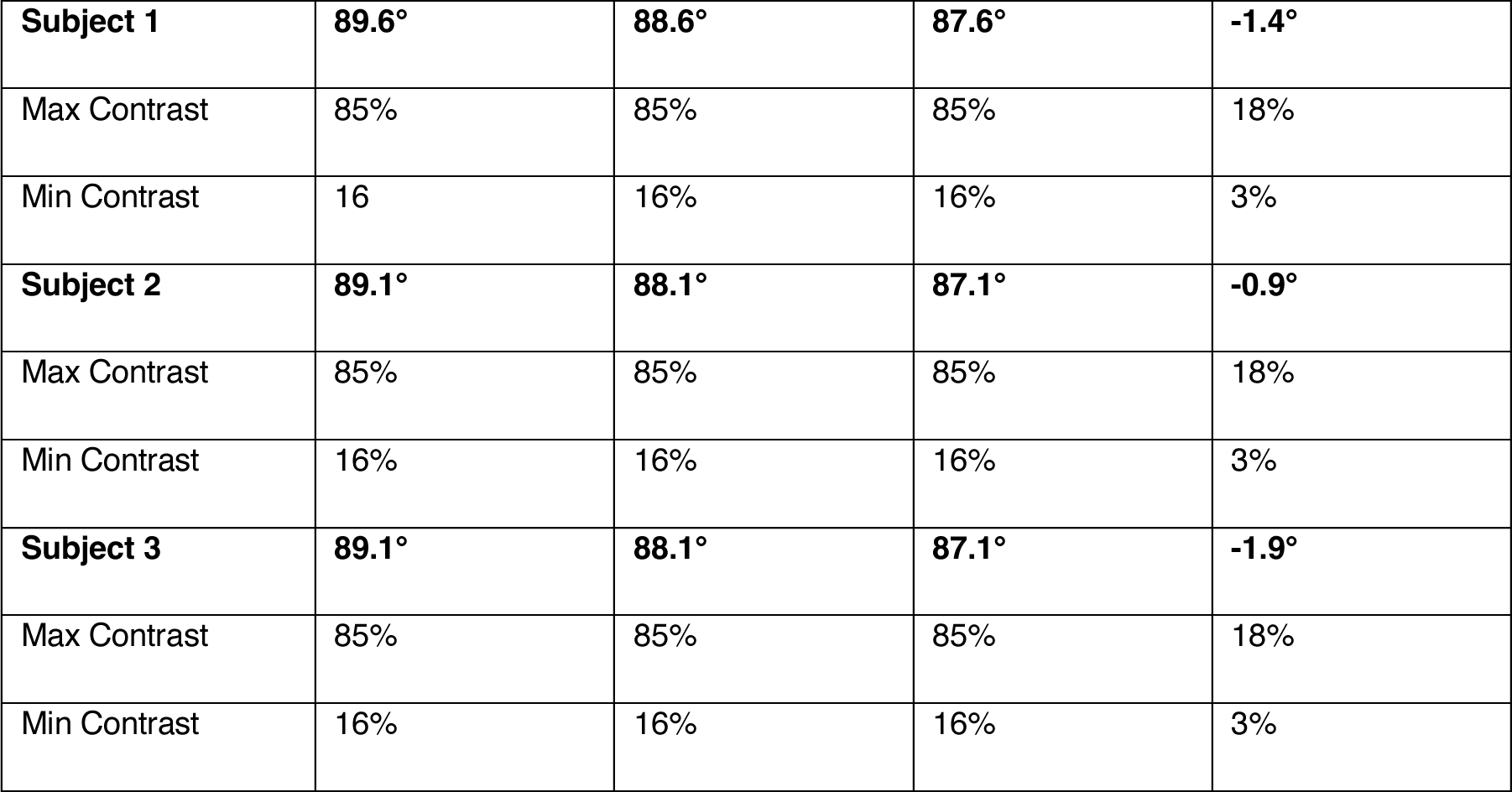
Subject Specific Directions and Contrasts – CTM. The chromatic directions and maximum/minimum contrasts of the tracking task. This table includes the chromatic directions specific to an individual subject.

|  |  |  |  |  |
| --- | --- | --- | --- | --- |
| **Subject 1** | **89.6°** | **88.6°** | **87.6°** | **-1.4°** |
| Max Contrast | 85% | 85% | 85% | 18% |
| Min Contrast | 16 | 16% | 16% | 3% |
| **Subject 2** | **89.1°** | **88.1°** | **87.1°** | **-0.9°** |
| Max Contrast | 85% | 85% | 85% | 18% |
| Min Contrast | 16% | 16% | 16% | 3% |
| **Subject 3** | **89.1°** | **88.1°** | **87.1°** | **-1.9°** |
| Max Contrast | 85% | 85% | 85% | 18% |
| Min Contrast | 16% | 16% | 16% | 3% |

**Table S3:** The Directions and Maximum Contrasts used in the Detection Task.

| Subject | -86.25° | -82.5° | -78.75° | -75° | -45° | 0° | 45° | 75° | 78.75° | 82.5° | 86.25° | 90° |
| --- | --- | --- | --- | --- | --- | --- | --- | --- | --- | --- | --- | --- |
| 1 | 14% | 10% | 9% | 6% | 2.5% | 1.75% | 2% | 7% | 7% | 10% | 18% | 18% |
| 2 | 10% | 7% | 5% | 4% | 1.25% | 1% | 1.25% | 4% | 5% | 7% | 10% | 9% |
| 3 | 10% | 8% | 5.5% | 6% | 1.5% | 1% | 1.5% | 4% | 5.5% | 8% | 10% | 15% |

## Footnotes

1 The orientation constrains the possible underlying mechanisms, it does not uniquely determine them. In particular, any given quadratic isoresponse ellipse is consistent with an equivalent contrast computed as the vector length of the joint response of many pairs of post-receptoral chromatic mechanisms (Poirson, Wandell, Varner, & Brainard, 1990; see also Barnett, Aguirre, & Brainard, 2021). None-the-less, the implication does hold that if two tasks are mediated by the same underlying mechanisms, possibly with differential sensitivity to the output of each mechanism, then the contours should have the same orientation.

